# A Pro-Regenerative Petroleum Jelly-Based, Copper-Doped Bioactive Glass Ointment for Impaired Wound Healing in Metabolic Syndrome

**DOI:** 10.64898/2026.06.12.731996

**Authors:** Huifeng Wang, Ophelia Tong, Youssef Ibrahim, Muskan Aslam, Yugang Liu, Chongwen Duan, Ruyue Luo, Angel Guo, Emma Vinokour, Andrew Kang, Pritika Jakka, Bin Jiang, Guillermo A. Ameer

**Author notes:** Corresponding author: Dr. Guillermo Ameer.

## Abstract

Chronic wound healing is often impaired in conditions such as metabolic syndrome, requiring effective therapeutic interventions to promote tissue regeneration and repair. In this study, we evaluated the wound healing potential of petroleum jelly (P Jelly)-based bioactive glass ointments (PBGCu) with varying copper concentrations (0, 1, and 3 wt%) in both *in vitro* and *in vivo* models of wound healing. PBGCu formulations demonstrated high biocompatibility with human dermal fibroblasts (HDF) and human umbilical vein endothelial cells (HUVEC). Additionally, PBGCu ointments exhibited strong antibacterial activity against *Staphylococcus aureus*, suggesting their utility for the care of chronic wounds. In both metabolic syndrome mouse and pig models, PBGCu3-treated wounds showed significantly faster wound closure, enhanced epithelial regeneration, and increased dermal thickness compared to saline and P Jelly controls. Histological analysis also revealed 50% increased vascularization (p < 0.0001) and a 90% reduction in scar formation (p < 0.0001) in PBGCu3-treated wounds. These findings show that PBGCu formulations, especially at 3 wt% copper concentration, significantly improve wound healing by promoting epithelial regeneration, dermal tissue formation, and vascularization, while also offering antibacterial protection. The sustained Cu^2+^ ions release from PBGCu ointments provides long-term support for tissue regeneration, positioning this ointment composition as a promising therapeutic tool for chronic wound management. Future studies will focus on elucidating the underlying mechanisms and evaluating the therapeutic efficacy of PBGCu formulations in infected wounds.

**Highlights:**

- Developed a Petroleum Jelly–based copper-doped bioactive glass ointment (PBGCu) enabling sustained and controlled Cu²⁺ ion release.
- PBGCu significantly accelerated wound closure and improved epithelial and dermal tissue regeneration.
- PBGCu enhanced hair follicle regeneration and tissue remodeling in full-thickness wounds.
- Validated therapeutic efficacy in both mouse and pig models that support translational relevance.
- Offers a simple, low-cost, and clinically adaptable topical formulation for metabolic syndrome-related wound complications.

## 1. Introduction

Metabolic syndrome (MetS), a cluster of conditions including central obesity, dyslipidemia, hypertension, and insulin resistance, has emerged as a major global health issue with far-reaching medical and economic ramifications.[1–3] Individuals with MetS often experience wound-healing complications stemming from a state of chronic low-grade inflammation and impaired microcirculation.[4, 5] These disruptions compromise essential phases of the wound-healing process, including hemostasis, inflammation, proliferation, and remodeling.[6, 7] As a result, skin injuries in MetS patients can exhibit delayed closure, higher infection rates, and poor tissue regeneration.[8] Thus, there is an urgent need for advanced therapeutic strategies that can simultaneously enhance tissue repair and mitigate the underlying challenges posed by MetS.

Bioactive glasses (BGs) have garnered significant interest as potent regenerative materials in both bone and soft tissue applications.[9–11] Owing to their capacity to release therapeutic ions (e.g., silicon, calcium, and phosphorus), BGs can stimulate cellular proliferation, promote neovascularization, and support the formation of new tissue matrices.[12, 13] Recent efforts to further refine the functional properties of BGs have focused on doping these materials with biologically active metal ions.[14, 15] Copper (Cu) ion, in particular, has received notable attention because it facilitates angiogenesis, provides antibacterial activity, and modulates cellular signaling pathways vital for efficient wound closure.[16–18] These attributes highlight copper-doped bioactive glass (BGCu) as a promising material to address complex wound-healing deficits present due to MetS.

To harness and deliver the benefits of BGCu in a clinically relevant manner, we developed a Petroleum Jelly (P Jelly)-based copper-doped bioactive glass ointment (PBGCu). P Jelly serves as a readily available and commonly used protective carrier that maintains a moist wound environment that is critical for optimal cell migration and matrix formation. Simultaneously, the PBGCu provides controlled release of Cu^2+^ ions, which can foster angiogenesis, support hair follicle regeneration, and exhibit antibacterial effects.[19] This dual strategy is particularly important in wound sites compromised by MetS, where factors such as persistent inflammation, diminished blood flow, and susceptibility to infection collectively hinder robust tissue repair.

In this work, we first synthesized and characterized BGCu powder using scanning electron microscopy (SEM) and energy-dispersive X-ray spectroscopy (EDX) to confirm its morphology and elemental composition. We then formulated the BGCu powder into a P Jelly ointment, and systematically measured the Cu^2+^ ion release profile via inductively coupled plasma mass spectrometry (ICP-MS). The antibacterial and tubulogenic properties of PBGCu were examined *in vitro* to evaluate its potential in both infection control and vascularization. Finally, we assessed the therapeutic efficacy of this ointment in two preclinical MetS models of full thickness skin excision wound: a mouse model and a pig model. We show that PBGCu treatment significantly improved wound closure rates, enhanced hair follicle regeneration, promoted tissue remodeling, and facilitated angiogenesis. Taken together, our findings underscore the potential of PBGCu as a topical therapy with the capacity to address the multifaceted challenges of impaired wound healing in MetS. By demonstrating its efficacy in both small- and large-animal models, we aim to advance PBGCu ointment as a next-generation wound care intervention suitable for clinical translation.

## 2. Materials and Methods

### 2.1 Materials

Tetraethyl orthosilicate (TEOS), triethyl phosphate (TEP), nitric acid, calcium nitrate tetrahydrate, copper oxide, and ammonia solution were purchased from Millipore Sigma (Burlington, MA) and used without further purification. Human dermal fibroblasts (HDF) and human primary umbilical vein endothelial cells (HUVEC) and all the cell culture media and their growth kits were purchased from ATCC (Manassas, VA).

### 2.2 Fabrication of copper doped bioactive glass (BGCu)

The copper doped bioactive glasses (BGCu) with varying copper compositions were prepared via an alkali-catalyzed sol-gel method as previously reported.[20] Briefly, a solution comprising 21.6 ml of TEOS and 2.8 ml of 2M nitric acid was dissolved in 13.9 ml of ultrapure water and 50 ml of ethanol. The resulting acid silica solution was mixed with 2.2 ml of TEP. Then, 14.04 g of calcium nitrate tetrahydrate and different amounts of copper oxide (0.367 g for BGCu1 and 1.1 g for BGCu3, 1 wt% and 3 wt% respectively) were added to the mixture and stirred until dissolved completely. While vigorously stirring, 10 ml of 1M ammonium solution was added to the solution. The resulting gel was kept in an oven at 60°C for 24 hours to eliminate residual water and ethanol and then heated at 600°C for 2 hours to yield copper-doped bioactive glass (BGCu).

### 2.3 Characterization of BGCu

The composition of BGCu was characterized by SEM and EDX. The Hitachi S4800 SEM instrument in Northwestern University Atomic and Nanoscale Characterization Experimental Center (NUANCE) was used to take SEM pictures and Hitachi SU8030 cFEG instrument was used for EDX analysis. BGCu was ground into fine particles and gently picked up onto carbon tape on stubs. The samples were coated with 10 nm of Osmium (Filgen OPC60A Osmium Coater) and SEM images were taken with accelerating voltage at 2000V. EDX was used to perform elemental analysis on each sample, and maps were collected for Ca, Si, O, P, C, and Cu for BG samples doped with copper. The particle sizes were analyzed using ImageJ software.

### 2.4 Cytotoxicity study of BGCu

Human dermal fibroblasts (HDF) were cultured in a humidified incubator, maintained at 37°C with 5% CO_2_, using fibroblasts growth medium. Extracts were obtained by immersing 50 mg/mL of BGCu in the culture medium, and the resulting extracts were collected after 72 hours at 37°C. Next, HDF cells were seeded in 96-well tissue culture plates at a density of 1 × 10^4^ cells/well and incubated overnight. Afterward, 100 µL of the extract and its serial dilutions (0.5, 2.5, 5, 10, and 25 mg/mL) were added to each well, and the cells were further incubated for 24 hours. The cell viability of the extract was then assessed using the MTT assay.

### 2.5 Fabrication of P Jelly copper-doped bioactive glass (PBGCU) ointments

The ointments containing P Jelly-BG were created through the combination of melted P Jelly and BGCu powder (25 wt%) with vigorous stirring at 80°C for 30 minutes. After cooling to room temperature, the mixture transformed into PBGCU ointment. The proportion of BGCu used was based on the cell viability of BGCu.

### 2.6 Copper ion release from PBGCU ointments

To quantify the copper ion release from PBGCu ointment formulations (PBGCu0, PBGCu1, and PBGCu3), 100 mg of each ointment was dissolved in 1 mL of phosphate-buffered saline (PBS). The solution was collected at different time points (1, 3, 5, 7, 9, and 12 days). To measure copper ion from the ointments, samples were first mixed with 1% nitric acid. After vortexing and sonication, the samples were incubated for one hour and then filtered to remove any particulate matter. The filtrates were subsequently diluted with deionized water and analyzed using inductively coupled plasma mass spectrometry (ICP-MS) to determine the copper concentration. Copper standards (0.1 µg/L, 1 µg/L, 10 µg/L, 100 µg/L, and 1 mg/L of copper nitrate) were prepared in the same nitric acid matrix to match the sample conditions, and the ICP-MS instrument was calibrated accordingly.

### 2.7 Antibacterial assay

Overnight culture of *S. aureus* was conducted in Luria-Bertani (LB) broth at 200 rpm and 37°C. A 48-well plate was prepared by adding 100 µL of PBS and 10 mg of PBGCu0, PBGCu1 and PBGCu3 ointments to each well and mixing it with 100 µL of bacteria suspension (in PBS, 1.5×10^4^ CFU/mL). PBS was used as a control. The plate was then incubated at 37°C for 24 h. The bacteria suspension was diluted with 800 µL of sterilized PBS, and 20 µL of the collected bacteria suspension was spread on an LB agar plate (15 mL of LB agar solution in a 10 mm Petri dish). The agar plates were incubated overnight, and digital images were taken. The colony-forming units (CFU) on the Petri dish were counted using ImageJ software. Each group was tested in triplicate, and the killing ratios of bacteria were calculated using the following equation:

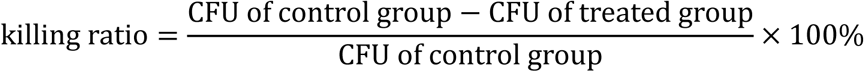

### 2.8 Cytotoxicity study of PBGCu ointments

To evaluate the cell compatibility of PBGCu ointments, a 24-well tissue culture plate was seeded with HDF or HUVEC at a density of 5 × 10^4^ cells per well in 24-well plates and incubated overnight at 37 °C. After that, the cells were exposed to 10 mg/mL PBGCu ointments for 24 hours. The cell viability was determined using alamarBlue assay and visualized by Live/Dead assay.

### 2.9 Cell migration assay

Scratch assays were conducted on HDF by creating a wound on the monolayer of cells. Subsequently, the cells were incubated at 37°C with different agents including saline, P Jelly, PBGCu0, PBGCu1 and PBGCu3 ointments. Digital images were captured at different time points. The wound area was quantified using ImageJ software by tracing the edge of the wounds, and the percentage of wound area was calculated using the following equation.

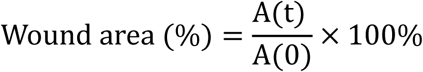

where A(t) is the wound area at time t and A(0) is its initial area.

### 2.10 Tubulogenesis assay

HUVECs were seeded in 24-well plates at a density of 5 × 10^4^ cells/well and left to incubate overnight. Afterward, the cells were exposed to different agents, including saline, P Jelly, PBGCu0, PBGCu1 and PBGCu3 ointments for 24 hours. Matrigel (BD Biosciences, San Jose, CA, USA) was diluted 2-fold with vascular basal medium and added to a 96-well plate at a volume of 60 μL per well. The plate was then pre-incubated at 37 °C for 30 minutes. The cells were collected and plated on the 96-well plate at a density of 2.5 × 10^4^ cells/well and then incubated for 4 hours. The resulting enclosed networks of complete tubes were digitally imaged and quantified using the Angiogenesis Analyzer plugin on ImageJ.

### 2.11 Evaluation of PBGCu ointment in a diabetic mouse splinted excisional wound model

The *in vivo* experiment to evaluate the wound healing performance of PBGCu ointment was initially conducted on male db/db mice (BKS.Cg-Dock7m +/+ Leprdb/J Homozygous for Leprdb) that were between 8 to 12 weeks old. All the mice had an average weight of approximately 50 g and blood glucose levels above 400 mg/dL. These mice were purchased from the Jackson Laboratory and were housed in the Center for Comparative Medicine at Northwestern University. All animal protocols were approved by Northwestern University’s Institutional Animal Care and Use Committee (IS000018748). To assess the effectiveness of the treatments, we used a splinted excisional wound model on the dorsal side of the mice, where a 6-mm circular full-thickness wound was created in the center of each splinted area. The mice were divided into 5 groups, each containing five animals, and each group was treated with one of the following: (1) PBGCu0, (2) PBGCu1, (3) PBGCu3 and (4) P Jelly and (5) saline as controls. To prevent skin contraction, doughnut-shaped silicon rubber splints were attached to both sides of the wound after depilation.[21] The wounds were treated every 3 days with 40 mg of PBGCu ointment or P Jelly or 40 µL of saline solution. A transparent sterile occlusive dressing TegaDerm^TM^ was placed over the wound and splint, and the wound dressings were changed every three days for a total of three times. Digital images of the wound area were taken every three days and quantified using ImageJ by normalizing the wound area to the known splint area at each time point.

### 2.12 Evaluation of PBGCu ointment in a metabolic syndrome swine excisional wound model

Ossabaw pigs with metabolic syndrome were obtained from Corvus Biomedical LLC and housed at the Center for Comparative Medicine at Northwestern University. All animal procedures were approved by Northwestern University’s Institutional Animal Care and Use Committee (IS000018748). To assess the efficacy of the treatments, eight full-thickness square wounds (3×3×2 cm^3^) were created on each animal, with four wounds on each side of the pigs’ backs. Three pigs were used in this study, and each pig received eight full-thickness wounds. The wounds were randomly assigned to four treatment groups: (1) PBGCu3, (2) PBGCu0, (3) P Jelly, and (4) saline, with each treatment applied to two wounds per pig. Each wound was treated with 2.5 g of PBGCu ointment or P Jelly, or 2.5 mL of saline solution. Wounds were treated once per week for 3 weeks and after each treatment they were covered with a transparent sterile occlusive dressing (TegaDerm™). Digital images of the wound area were taken weekly and analyzed using ImageJ software to quantify wound closure at each time point.

### 2.13 Tissue processing

Upon complete wound closure (27 days post-wounding for mice and 8 weeks for pigs), the animals were euthanized, and the regenerated wound tissue was collected. For mice, tissue was excised using a 10-mm biopsy punch (Acuderm, Fort Lauderdale, FL). For pigs, a tissue cube (3×3×2 cm³) was cut. The tissue was fixed in 4% paraformaldehyde overnight, washed with PBS twice every 2 hours and then overnight. Following fixation, the tissue was dehydrated using gradually ascending ethanol solutions (70%, 80%, 95%, and 100% twice) for 45 minutes each. The tissue was then cleared using xylene twice for 1 hour each time, and paraffin overnight at 60°C. The next day, the tissue was transferred to an embedding machine and immersed in paraffin overnight before being embedded in a paraffin mold.

### 2.14 Histological analysis

The tissues were sectioned (5-µm thickness) and placed onto microscope slides. Tissue sections were then stained for hematoxylin and eosin (H&E) and Masson’s trichrome staining (MTS) for both mouse and pig tissues. The slides were then imaged using Hamamatsue NanoZoomer 2.0 HT (Hamamatsu, Japan). The epidermal and dermal thickness and scar length were quantified using NDP view2 software.

### 2.15 Statistical analysis

All data are presented as mean ± standard deviation (SD) (n = 3 for all *in vitro* studies and n = 6 for all *in vivo* studies). Statistical significance was assessed using one-way ANOVA with Bonferroni multiple comparison corrections. Statistical analyses were conducted using GraphPad Prism 9 software, with significance levels indicated as *p < 0.05, **p < 0.01, ***p < 0.001, and ****p < 0.0001.

## 3. Results

### 3.1 BGCu and petroleum jelly can be formulated to enable sustained release of Cu^2+^ ions

The bioactive glass powders were successfully fabricated using a sol-gel process, followed by aging. These powders were then mixed with melted P Jelly at 80 °C for 30 min to create the PBGCu ointments, with copper concentrations of 0, 1, and 3 wt% (**Figure 1A**). To determine the concentration of copper oxide used in the PBGCu ointments, we first performed a cell viability assay using extracts from BGCu powder (50 mg/mL) incubated at 37°C for 72 hours. The extracts and their serial dilutions (0.5, 2.5, 5, 10, and 25 mg/mL) were then added to L929 fibroblasts, and cell viability was assessed using the MTT assay. Cells cultured with 25 mg/mL of all bioactive glass formulations maintained over 80% viability (**Figure S1**). Based on these results, we mixed 25% BGCu with 75% P Jelly to prepare the ointment. The digital images of BGCu powders (**Figure 1B**) and PBGCu ointments (**Figure 1C**) demonstrate that increasing copper concentration leads to a gradual change in the appearance of the powders and ointments. The powders range from white (BGCu0) to darker shades of grey to black with increasing copper content (BGCu3). Similarly, the ointments show a shift in color, with increasing copper concentration influencing the ointment’s visual properties. SEM images revealed that the BGCu powders consisted of fine particles with an average particle size of 56.2 ± 20.2 µm. EDX analysis demonstrated a uniform distribution of key elements in the BGCu powders, including silicon, calcium, phosphorus, and copper (**Figure 1D**). Copper is clearly detected in BGCu1 and BGCu3, with the intensity of copper signal increasing as copper content increases (1.09% and 3.02%). The Cu^2+^ ions release from PBGCu ointments was evaluated over a 12-day period using ICP-MS (**Figure 1E**). The data show a gradual increase in Cu^2+^ ions release over time, with ointments containing 3 wt% copper demonstrating higher release rates compared to the 1 wt% copper ointment. This indicates that the PBGCu ointments can serve as a potential vehicle for controlled copper delivery. Overall, these results highlight the successful incorporation of copper into bioactive glass powders and ointments, with potential applications in controlled metal ion release for biomedical uses.

**Figure 1:**
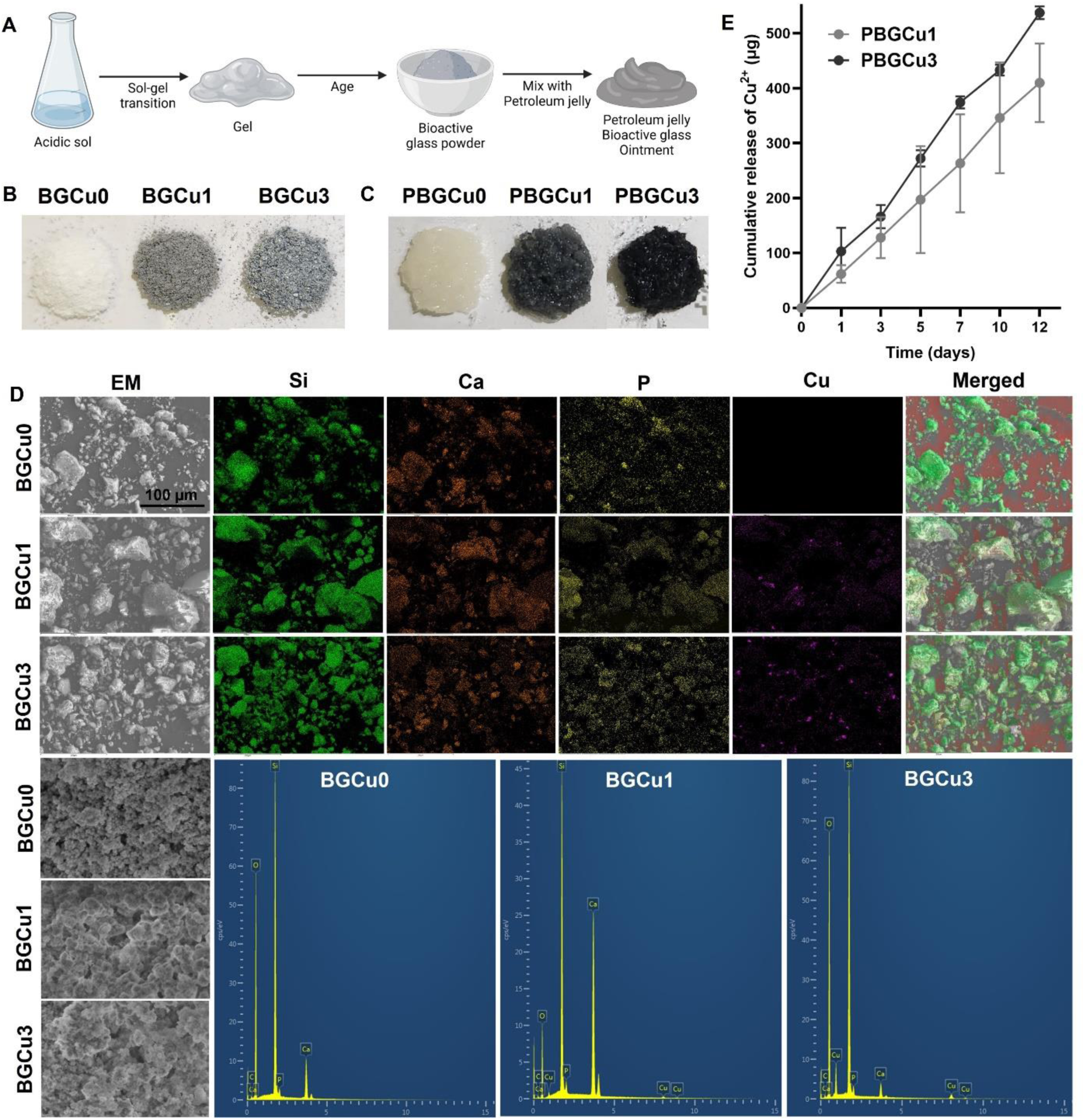
Fabrication and characterization of bioactive glass (BGCu) powders and P Jelly-based bioactive glass ointments (PBGCu) with varying copper concentrations. (A) Schematic of the fabrication process for BGCu powder and PBGCu ointment, including the sol-gel transition, aging, and mixing with P Jelly. (B) Digital images of BGCu powders with different copper concentrations (0, 1, and 3 wt%). (C) Digital images of PBGCu ointments with different copper concentrations showing the visual differences with increasing copper ions content. (D) SEM images and EDX analysis of BGCu powders, with elemental mappings for silicon (Si), calcium (Ca), phosphorus (P), and copper (Cu), along with the merged image highlighting copper distribution. (E) Cumulative Cu^2+^ ion release from PBGCu ointments over 12 days, showing a gradual increase in copper release, with ointments containing 1% and 3 wt% copper exhibiting higher release rates.

### 3.2 PBGCu is cell compatible and has antibacterial properties

Both human dermal fibroblasts (HDF) and human umbilical vein endothelial cells (HUVEC) exhibited high viability when cultured with PBGCu ointments at copper concentrations of 0 %, 1 %, and 3 wt%. Live/dead assays showed no significant cytotoxicity and effect on cell morphology. The cell viability assays confirmed that all PBGCu maintained cell viability comparable to the control groups, indicating that the ointments are biocompatible (**Figure 2A-D**). The antibacterial efficacy of the PBGCu ointments was evaluated using *Staphylococcus aureus* bacteria. While the control group exhibited extensive bacterial growth, the PBGCu formulations showed reduced colony formation, with the ointments containing copper being more effective at inhibiting bacterial growth (**Figure 2E**). Quantitative analysis of the killing ratio confirmed a significant antibacterial effect, with all formulations demonstrating over 80% efficacy (p<0.0001). Notably, the copper-containing formulations exhibited exceptional antibacterial activity, achieving a killing ratio exceeding 99% (p<0.0001) (**Figure 2F**).

**Figure 2:**
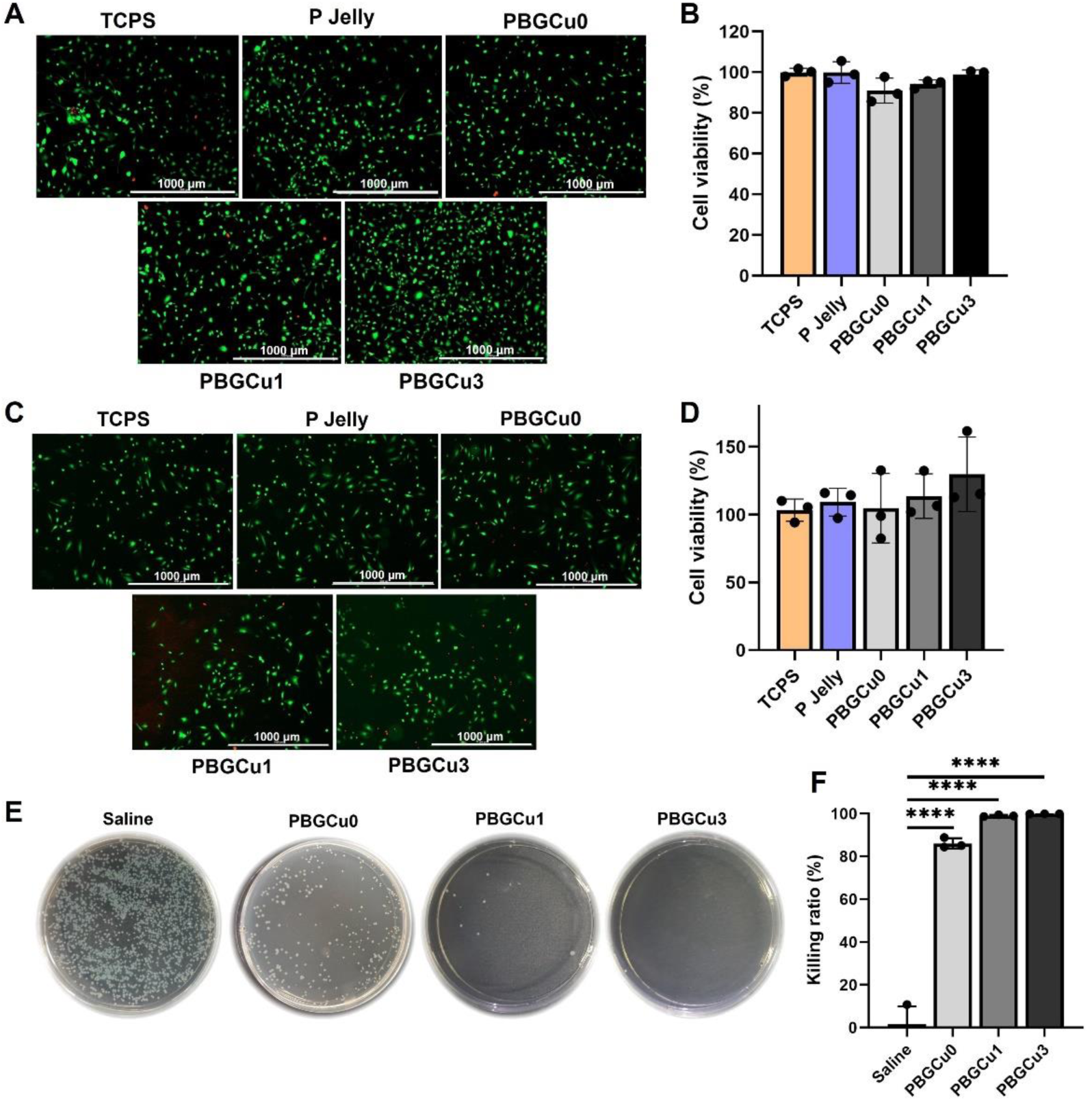
Biocompatibility and antibacterial properties of P Jelly-based bioactive glass ointments (PBGCu) with varying copper concentrations. **(A)** Live/dead staining images of human dermal fibroblasts (HDF) cultured with PBGCu ointments (0 %, 1 %, and 3 wt% copper). The scale bar is 1000 µm. **(B)** Cell viability of HDFs treated with PBGCu ointments. **(C)** Live/dead staining images of human umbilical vein endothelial cells (HUVEC) cultured with PBGCu ointments. The scale bar is 1000 µm. **(D)** Cell viability of HUVECs treated with PBGCu ointments. **(E)** Bacterial colony growth of Staphylococcus aureus after treatment with PBGCu ointments. **(F)** Quantification of the antibacterial killing ratio. All data are presented as mean ± standard deviation (SD), and statistical analysis was performed using one-way ANOVA followed by Bonferroni post hoc correction. (*n* = 3, **p < 0.01, ***p < 0.001, ****p < 0.0001).

### 3.3 PBGCu promotes HDF migration and HUVEC tubulogenesis *in vitro*

The HDF migration assay showed that the application of PBGCu ointments enhanced cell migration, with a greater reduction in wound area observed in ointments containing copper (**Figure 3A**). The 1% and 3 wt% copper formulations significantly enhanced migration compared to TCPS, with a 25% and 37% reduction in wound area, respectively (p = 0.0033 and 0.0002). Furthermore, PBGCu3 demonstrated a 24% reduction in wound area compared to PBGCu0 (p = 0.018) (**Figure 3B**). The HUVEC tubulogenesis assay demonstrated that PBGCu3 facilitated the formation of more organized and interconnected tubular structures (**Figure 3C**). Quantification of tubule formation showed that PBGCu3 significantly increased the number of tubule nodes, junctions, and meshes, with a 66%, 64%, and 70% increase, respectively, compared to saline (p = 0.0284, 0.0435, and 0.0364). These results indicate enhanced angiogenesis and tissue network formation (**Figures 3D-F**). Overall, the results suggest that PBGCu ointments, especially those with higher copper concentrations, effectively promote both cell migration and tubulogenesis, indicating their potential as therapeutic agents for wound healing and tissue regeneration.

**Figure 3:**
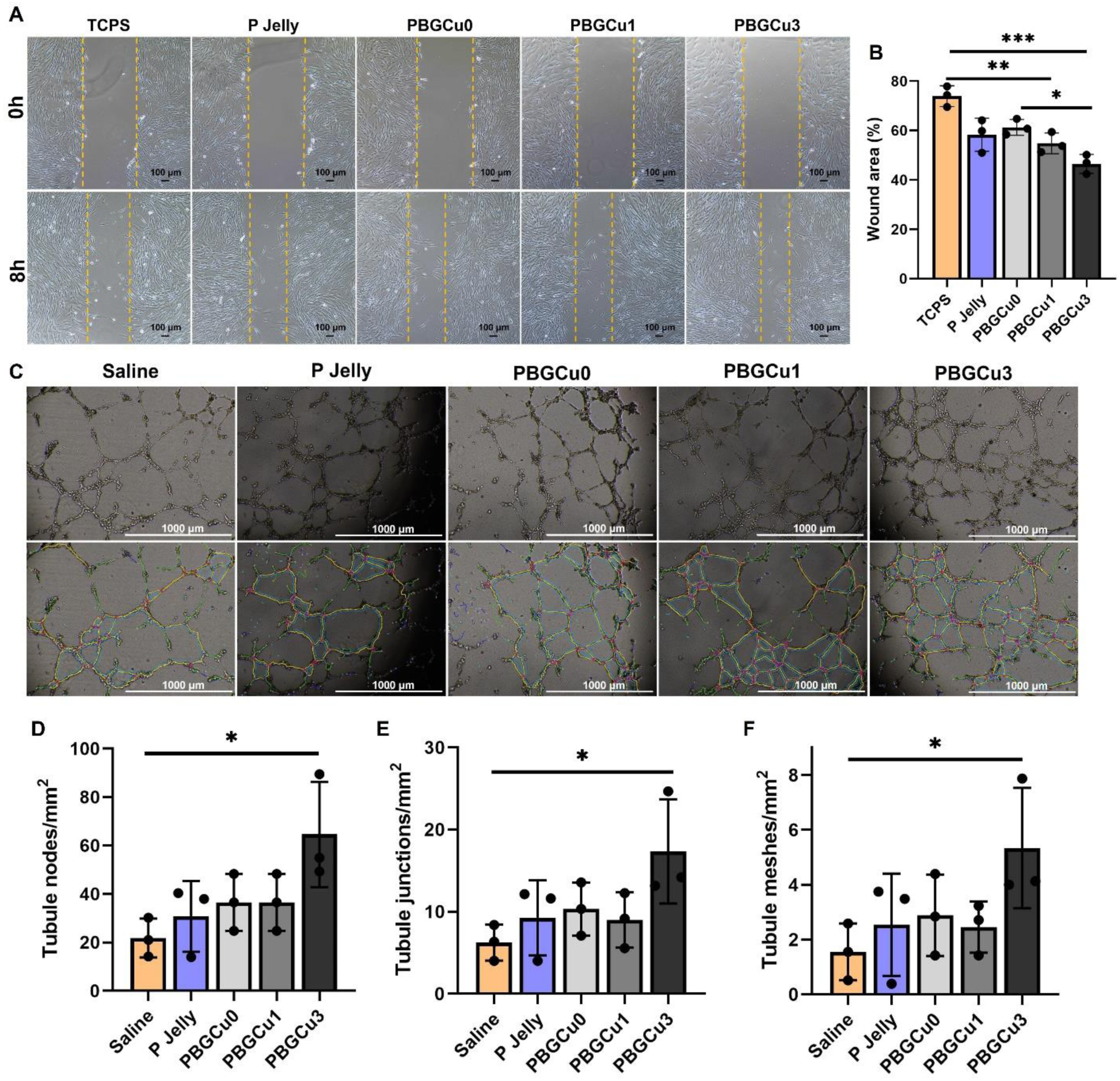
PBGCu ointments promote HDF migration and HUVEC tubulogenesis *in vitro*. (**A**) Representative images from the HDF migration assay at 0 hours and 8 hours after treatment with PBGCu ointments (0, 1, and 3 wt% copper). The scale bar is 100 µm. (**B**) Quantification of wound area. (**C**) Representative images from the HUVEC tubulogenesis assay after treatment with PBGCu ointments. The scale bar is 1000 µm. Quantification of tubule (**D**) nodes, (**E**) junctions, and (**F**) meshes in PBGCu ointments. All data are presented as mean ± standard deviation (SD), and statistical analysis was performed using one-way ANOVA followed by Bonferroni post hoc correction. (*n* = 3, **p < 0.01, ***p < 0.001, ****p < 0.0001).

### 3.4 PBGCu accelerates wound closure in diabetic mice with metabolic syndrome

Based on the *in vitro* findings, which demonstrated enhanced proliferation and migration of dermal fibroblasts, as well as the antibacterial and tubulogenic properties of PBGCu, we hypothesized that these factors would lead to faster wound closure rate *in vivo*. To test this hypothesis, we evaluated the efficacy of PBGCu as a regenerative dressing in a splinted excisional full-thickness wound model in *db/db* mice, a commonly used model for metabolic syndrome (**Figure 4A**).[22] The blood glucose levels of mice were constantly monitored throughout the experiment period. At all measured time points, blood glucose levels remained above 200 mg/dL in all mice (**Figure S2**). The splinted excisional wound model was chosen to minimize skin contraction, allowing for a more accurate assessment of tissue regeneration. The results demonstrated that wounds treated with PBGCu3 healed significantly faster, with a 61.5±12.7% reduction in wound area compared to saline and P Jelly-treated wounds, which showed reductions of 32.5±10.8% and 36.0±8.4%, respectively at day 9 (p = 0.0008 and 0.001). By day 18 post-wounding, wounds treated with PBGCu3 were completely closed, while saline-treated wounds still had 30% of the wound area remaining open (p < 0.0001) (**Figure 4B-C**).

**Figure 4.**
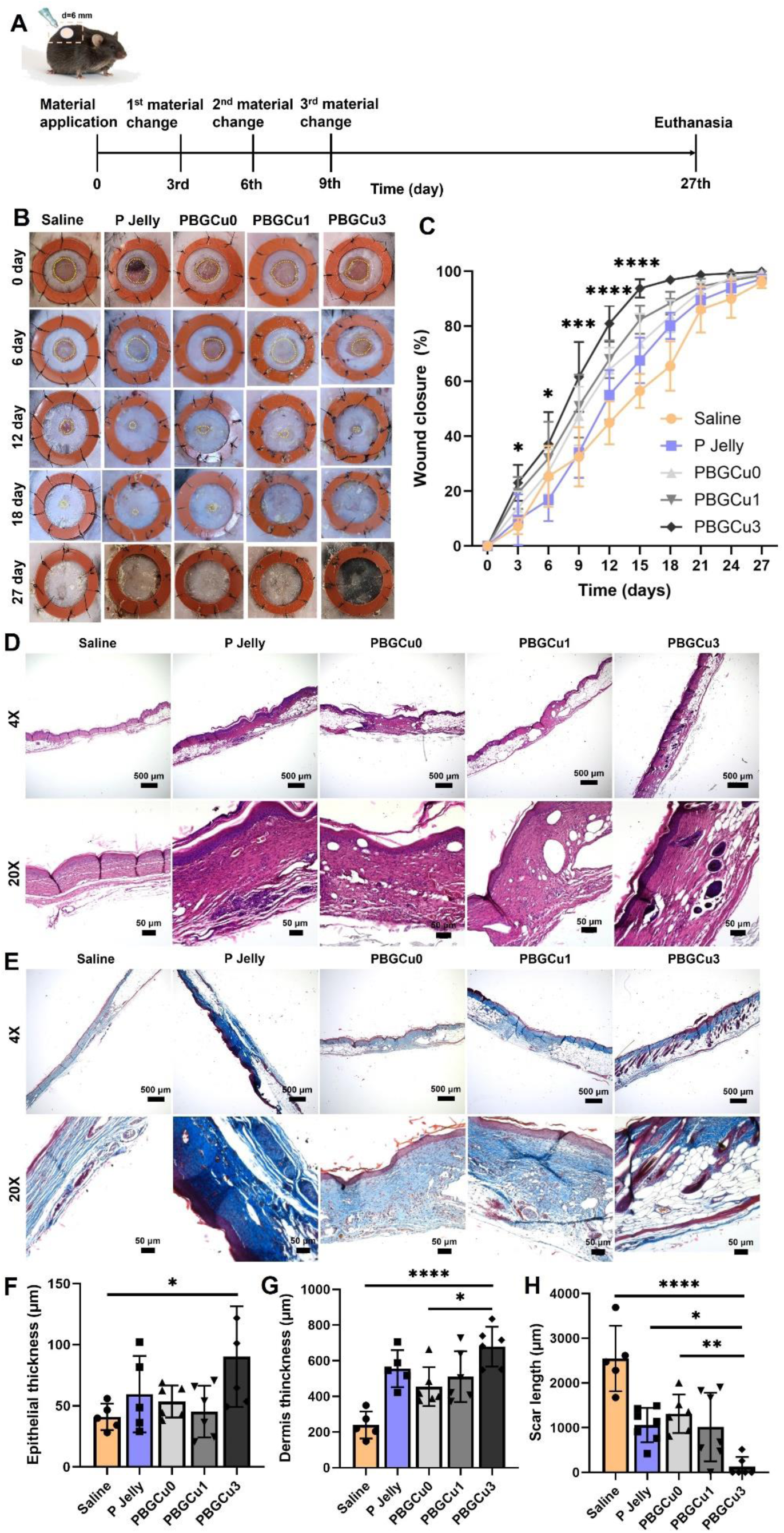
PBGCu promotes wound healing and tissue regeneration in a full-thickness excisional wound model in mice with diabetes and metabolic syndrome. (**A**) Experimental timeline outlining the application of materials (saline, P Jelly, and 0%, 1%, and 3% PBGCu formulations) at specified intervals, followed by euthanasia on day 27. (**B**) Representative images of wound closure at various time points (0, 6, 12, 18, and 27 days) for each treatment group. (**C**) Quantification of wound closure rates over time. (**D**) Hematoxylin and eosin (H&E) staining of tissue sections at 4x and 20x magnification. (**E**) Masson’s trichrome staining of tissue sections at 4x and 20x magnification. The scale bar for the 4x magnification is 500 µm. The scale bar for the 20x magnification is 50 µm. Quantification of (**F**) epithelial layer thickness, (**G**) dermal layer thickness, and (**H**) scar length. All data are presented as mean ± standard deviation (SD), and statistical analysis was performed using one-way ANOVA followed by Bonferroni post hoc correction. (*n* = 6, **p < 0.01, ***p < 0.001, ****p < 0.0001).

### 3.5 PBGCu promotes epithelial and dermal tissue formation as well as hair follicle regeneration

On day 27, regenerated skin tissue samples were collected, processed, and stained with hematoxylin and eosin (H&E) and Masson’s trichrome staining (MTS) for histological analysis (**Figure 4D** and **4E**). The epidermal layer of PBGCu3-treated wounds was 44 µm thicker than that of saline-treated wounds (p = 0.0374). Dermal regeneration in the wound beds was significantly enhanced in PBGCu3-treated wounds, which exhibited approximately 438.9 µm and 210.0 µm greater thickness compared to saline- and PBGCu0-treated wounds, respectively (p < 0.0001 and p = 0.0166) (**Figure 4F** and **4G**). Furthermore, hair follicle regeneration was markedly enhanced, and scar formation was significantly reduced in the PBGCu3-treated group compared with the saline-, P Jelly–, and PBGCu0-treated wounds. Specifically, the scar length in the PBGCu3-treated wounds was reduced by 95% (p < 0.0001), 88% (p = 0.037), and 90% (p = 0.0071), respectively (**Figure 4H**). These findings suggest that PBGCu3 not only accelerates wound closure but also promotes the regeneration of essential skin structures, such as the dermis, epithelium, and hair follicles, which are crucial for functional skin recovery.

### 3.6 PBGCu accelerates wound closure in pigs with metabolic syndrome

To enhance the translational relevance of our findings, we assessed the therapeutic efficacy of PBGCu ointment in a swine model of metabolic syndrome. The model was validated by elevated body weight, blood glucose, total cholesterol, and triglyceride levels, consistent with previously reported metabolic syndrome profiles (**Table S1**).[23] Three pigs were included in the study, each receiving eight treatments. Following surgical wound creation, PBGCu ointment was applied and replaced at weeks 1 and 2 (**Figure 5A**). Wound healing progression was monitored weekly through imaging and measurement until week 7, after which the animals were euthanized. Due to inter-animal variability in wound healing responses, wound closure curves were plotted individually for each pig. Treatment efficacy was therefore evaluated within each animal, comparing wound closure rates across treatment groups. The PBGCu3 formulation exhibited the most rapid wound closure, with a significant reduction in wound volume (77.1±7.9% in Pig 1, 39.4±7.8% in Pig 2 and 47.4±2.6% in Pig 3) compared to saline (53±4.2%, p=0.0495 in Pig 1, 9.1±4.2%, p=0.0467 in Pig 2 and 7.7±6.0%, p=0.0473 in Pig 3), P Jelly (57.7±5.4%, p=0.0935 in Pig 1, 13.6±12.1%, p=0.0841 in Pig 2 and 21.8±8.5%, p=0.1857 in Pig 3), and PBGCu0 (61.9±5.7%, p=0.1804 in Pig 1, 16.1±0.5%, p=0.1133 in Pig 2 and 15.9±6.3%, p=0.1077 in Pig 3) during the first week (Figure 5B,D, **S3A,C** and **S4A,C**). By week 2, PBGCu3-treated wounds showed a 81.6±3.3%, 60.9±7.5% and 90.6±0.8% reduction in wound area in Pig 1, 2 and 3, respectively, compared to saline (48.3±0.3%, p=0.006 in Pig 1, 36.1±10.3%, p=0.0475 in Pig 2 and 61.6±3.6%, p=0.014 in Pig 3), P Jelly (49.5±6.4% in Pig 1, p= 0.0069, 34.9±3.9%, p=0.0394 in Pig 2 and 67.5±6.7%, p= 0.0313 in Pig 3), and PBGCu0 (52.4±5.3% in Pig 1, p= 0.0097, 42.0±11.5%, p=0.1773 in Pig 2 and 61.3±6.2%, p= 0.0136 in Pig 3) treated wounds (**Figure 5C, S3C** and **S4C**). Over the course of the study, PBGCu-treated groups consistently outperformed the saline and P Jelly controls in terms of both wound closure and tissue regeneration. These results suggest that PBGCu accelerates wound closure and enhance tissue healing in pigs with metabolic syndrome, a condition typically associated with impaired healing responses. The accelerated wound healing observed in this model emphasizes the potential of PBGCu as a promising therapeutic option for improving wound healing in individuals with metabolic conditions.

**Figure 5.**
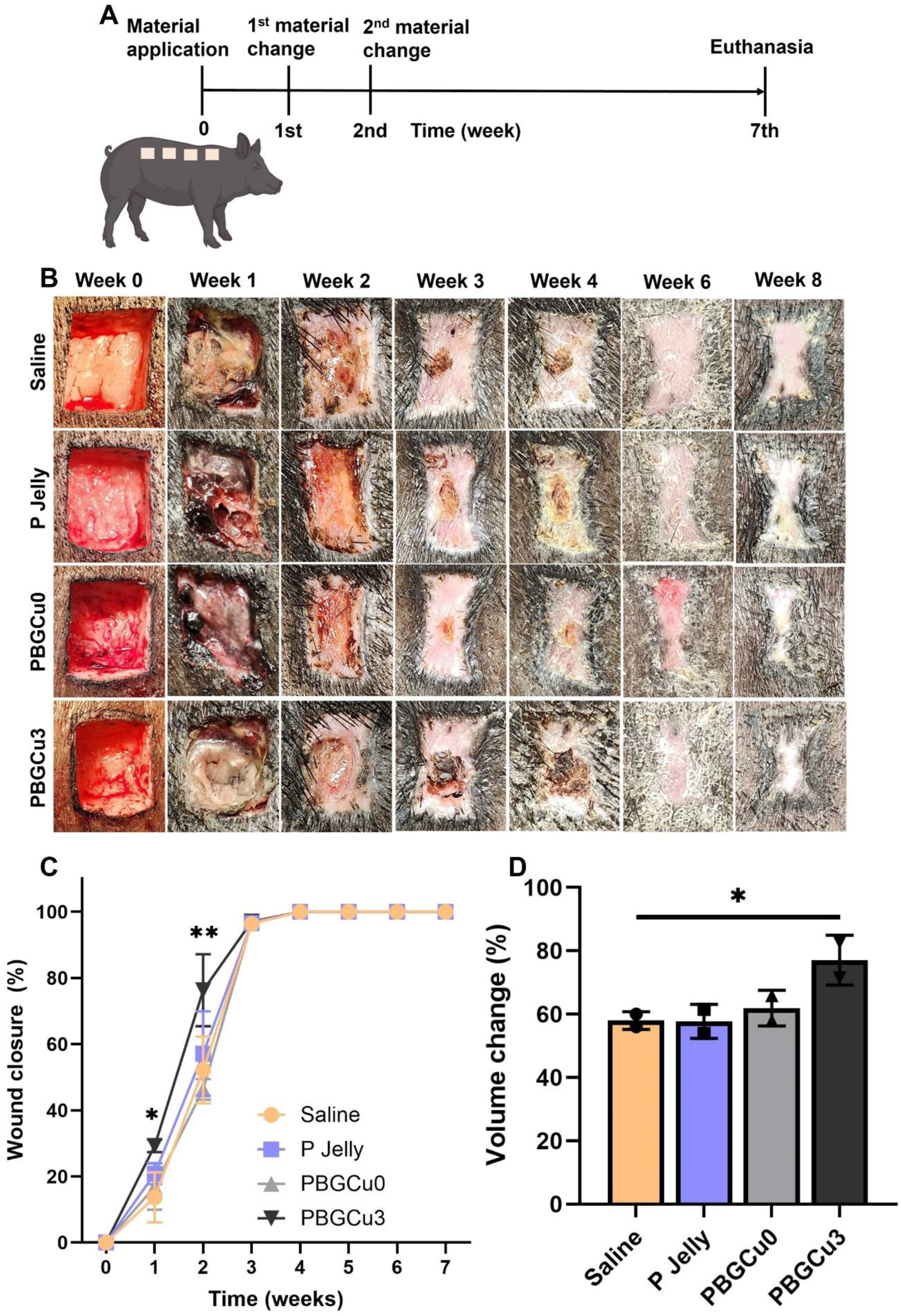
PBGCu accelerates wound healing of full-thickness excisional wound in pigs with metabolic syndrome. (**A**) Experimental timeline for the application of materials (saline, P Jelly, and 0% and 3% PBGCu formulations) at specified intervals, followed by euthanasia at week 7. (**B**) Representative images showing wound closure at various time points (1, 2, 3, 4, 6, and 8 weeks) for each treatment group in Pig 1. (**C**) Quantification of wound closure rates over time in Pig 1, with PBGCu-treated groups showing significantly faster healing, and PBGCu3 demonstrating the most rapid closure. (**D**) Quantification of wound volume change after 1 week in Pig 1, with significant differences observed between PBGCu-treated groups and controls. All data are presented as mean ± standard deviation (SD), and statistical analysis was performed using one-way ANOVA followed by Bonferroni post hoc correction. (*n* = 6, **p < 0.01, ***p < 0.001, ****p < 0.0001).

### 3.7 PBGCu promotes epithelial and granulation tissue formation and vascularization of full thickness skin wounds in pig

We conducted a histological analysis of regenerated skin tissue from pig wounds treated with saline, P Jelly, PBGCu0, and PBGCu3. PBGCu3-treated wounds demonstrated thicker and more well-structured epithelial layers, with the most substantial increase in epithelial thickness observed (**Figures 6A**-**B**). The epidermal layer of PBGCu3-treated wounds was 98.8 µm (p < 0.0001), 90.0 µm (p = 0.0002), and 91.7 µm (p = 0.0002) thicker than those of saline-, P Jelly-, and PBGCu0-treated wounds, respectively. Additionally, Masson’s trichrome staining highlighted enhanced collagen deposition and dermal regeneration in PBGCu3-treated wounds (**Figure 6C**). This was further supported by quantification of dermal thickness, where PBGCu3-treated wounds displayed increased dermal regeneration (**Figure 6D**). Specifically, dermal thickness in the wound beds was significantly greater in PBGCu3-treated wounds, with approximately 2.45 mm (p = 0.0411) and 2.56 mm (p = 0.0311) more thickness compared to saline- and P Jelly-treated wounds, respectively. Furthermore, blood vessel density was significantly higher in PBGCu3-treated wounds, showing increases of 51.7% (p < 0.0001), 47.3% (p = 0.0002), and 42.6% (p = 0.0005) compared to saline-, P Jelly-, and PBGCu0-treated wounds. This suggests improved vascularization, which is crucial for enhanced tissue repair (**Figures 6E-F**). Overall, these findings underscore the effectiveness of PBGCu in promoting wound healing by enhancing epithelial regeneration, dermal tissue formation, and vascularization.

**Figure 6.**
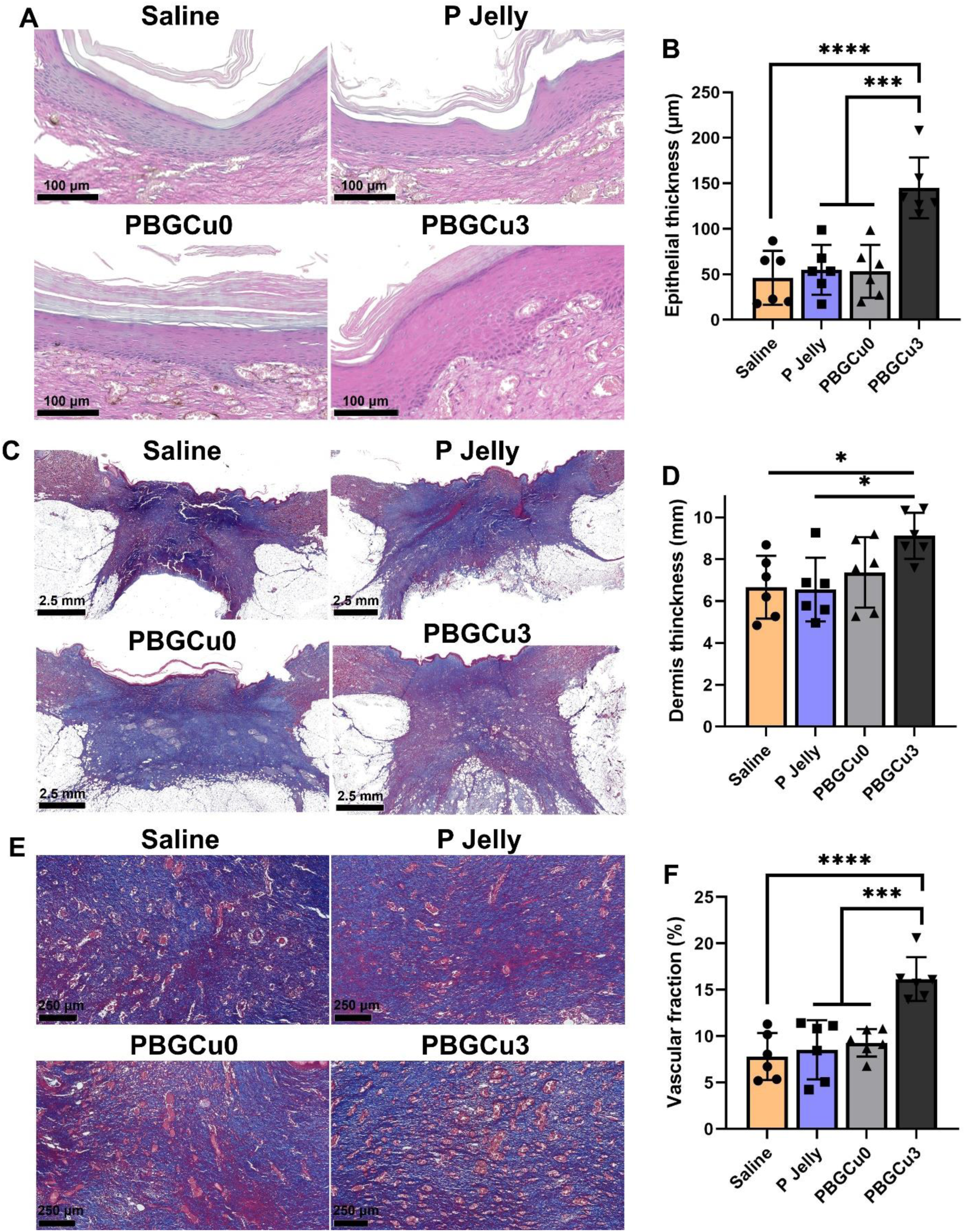
PBGCu promotes epithelial regeneration, dermal thickness, and vascularization in pig full thickness skin wounds. (**A**) H&E staining of tissue sections showing epithelial regeneration in wounds treated with saline, P Jelly, PBGCu0, and PBGCu3. The yellow arrows indicate the thickness of epithelial layer. The scale bar is 100 µm. (**B**) Quantification of epithelial thickness. (**C**) Masson’s trichrome staining of tissue sections. The yellow arrows indicate the thickness of dermal layer. The scale bar is 2.5 mm. (**D**) Quantification of granulation tissue formation. (**E**) Masson’s trichrome staining at higher magnification. The yellow triangles indicate blood vessels. The scale bar is 250 µm. (**F**) Quantification of vascular fraction. All data are presented as mean ± standard deviation (SD), and statistical analysis was performed using one-way ANOVA followed by Bonferroni post hoc correction. (*n* = 6, **p < 0.01, ***p < 0.001, ****p < 0.0001).

## 4. Discussion

The results of this study highlight the significant potential of PBGCu formulations, particularly PBGCu3, in promoting wound healing and tissue regeneration. Both *in vitro* and *in vivo* findings demonstrate that PBGCu formulations, which combine copper-doped bioactive glass with a petroleum jelly base for sustained Cu^2+^ ions release, provide therapeutic benefits for wound healing, especially in conditions with impaired healing, such as metabolic syndrome.

In our *in vitro* studies, PBGCu formulations exhibited high biocompatibility, with no significant cytotoxicity observed in human dermal fibroblasts and endothelial cells. These results suggest that PBGCu formulations are non-toxic to essential cell types involved in wound healing, a critical factor for their clinical application. Furthermore, PBGCu formulations demonstrated strong antibacterial activity, with the copper-containing groups exhibiting the most pronounced effects, particularly against Staphylococcus aureus, a common pathogen associated with chronic wounds. Notably, PBGCu0 also inhibited bacterial growth, likely due to the release of calcium ions, which is consistent with previous reports. [24] Our findings further demonstrate that PBGCu formulations can offer both effective antibacterial properties and biocompatibility, which are crucial for successful wound management.

To further validate their therapeutic potential, *in vivo* studies were performed using the *db/db* mouse model, a well-established model of metabolic syndrome. Previous studies have reported characteristic metabolic parameters in this model, including significantly increased food intake, plasma insulin levels, and body fat ratio.[25] The PBGCu3-treated wounds exhibited significantly faster wound closure compared to the saline and P Jelly control groups, which highlight the efficacy of PBGCu in accelerating wound healing in a model that typically exhibits delayed healing due to metabolic dysfunction. Histological analysis demonstrated enhanced epithelial regeneration and dermal tissue formation in PBGCu3-treated wounds, with significant increases in both epithelial and dermal thickness. These results indicate that PBGCu3 formulations stimulate the regeneration of both epithelial and dermal cells in the wound site, which are consistent with the previous studies.[26, 27]

Additionally, we observed a reduction in scar formation in PBGCu3-treated wounds. The ability of PBGCu formulations to both accelerate 30% wound closure and improve tissue remodeling is a significant advantage, as excessive scarring can lead to impaired skin integrity. Our study builds on these findings by showing that PBGCu formulations not only promote wound healing but also stimulate hair follicle regeneration, making them particularly beneficial for achieving optimal functional outcomes in clinical applications.

We next evaluated the therapeutic efficacy of PBGCu in a clinically relevant metabolic syndrome (MetS) Ossabaw pig model, which closely recapitulates key features of human Mets including elevated body fat ratio, total cholesterol, and triglyceride concentrations (**Table S1**).[28] While copper-doped biomaterials have been investigated in small animal models, their evaluation in large animal MetS pig models remains limited. In this context, PBGCu3-treated wounds exhibited significantly faster wound closure when compared to control wounds. Histological analysis further revealed enhanced epithelial layer formation and granulation tissue, both of which are indicative of improved wound healing. More importantly, PBGCu3-treated wounds showed increased vascularization, with higher blood vessel density observed in the dermis. Vascularization is a critical component of wound healing, as it ensures an adequate supply of oxygen, nutrients, and immune cells to the regenerating tissue.[29] In chronic wounds or metabolic disorders, vascular insufficiency often hinders healing, and the ability of PBGCu to stimulate angiogenesis is a crucial benefit in such environments.[18] The enhanced vascularization observed in PBGCu3-treated wounds suggests that copper plays a key role in promoting angiogenesis, thereby contributing to faster and more effective wound healing.

Overall, the findings from this study suggest that PBGCu formulations, especially with 3 wt% copper, can significantly improve wound healing by promoting epithelial regeneration, dermal tissue formation, and vascularization. These formulations also exhibit strong antibacterial properties, making them an attractive option for chronic wound care, particularly in conditions like metabolic syndrome, where wound healing is often compromised. The sustained Cu^2+^ ions release from PBGCu ointments offers an advantage over other wound healing treatments by providing long-term support for tissue regeneration and reducing the frequency of treatment application.

Compared with existing studies that have explored the role of copper in promoting wound healing, our work demonstrates that PBGCu formulations uniquely integrate sustained copper release from bioactive glass and a petroleum jelly matrix. Prior reports have shown that copper ions can promote angiogenesis, antibacterial activity, and tissue regeneration; however, many of these systems, such as copper-loaded dressings or nanoparticle-based formulations, typically exhibit rapid or burst release profiles, with the majority of Cu^2+^ ions released within hours to the first 1-2 days. [30, 31] In contrast, our PBGCu formulation demonstrated a sustained Cu²⁺ release over an extended period (up to ∼12 days), enabling prolonged biological activity at the wound site. This sustained release translated into effective therapeutic outcomes, including strong antibacterial activity against Staphylococcus aureus and enhanced angiogenesis, as evidenced by increased tubulogenesis in vitro and higher vessel density *in vivo*. Compared to previously reported copper-based systems, which often show transient antibacterial effects or limited long-term vascularization, PBGCu provides continuous ion delivery that supports both infection control and tissue regeneration throughout the healing process. Furthermore, PBGCu treatment resulted in accelerated wound closure and improved tissue remodeling in both mouse and pig models, underscoring its potential advantage over conventional copper-based dressings that rely on short-term ion availability.[32] Additionally, The incorporation of copper-doped bioactive glass into a petroleum jelly base provides both biological and practical advantages. P Jelly is widely used in clinical wound care due to its ability to maintain a moist wound environment, which is essential for re-epithelialization, prevention of scar formation and protection against external contamination. It also serves as an occlusive barrier that reduces transepidermal water loss and support cell migration.[33] Commercial products such as Aquaphor and Vaseline-based formulations are commonly used for minor wound care and skin protection, highlighting the clinical relevance and safety of this carrier system. In this study, the combination of P Jelly with copper-doped bioactive glass enables not only effective topical delivery but also sustained release of bioactive Cu^2+^ ions. This synergistic design enhances bioactivity by promoting angiogenesis, collagen deposition, and tissue regeneration, ultimately leading to improved wound healing outcomes. [19] In contrast, many conventional copper-based biomaterials, such as nanoparticles, do not provide sustained release and may induce cytotoxicity, which limits their long-term effectiveness in chronic wound care.

Previous studies have demonstrated that copper-mediated wound healing is associated with the activation of pro-angiogenic pathways (e.g., HIF-1α/VEGF signaling), modulation of inflammatory responses, and stimulation of fibroblast proliferation and collagen deposition.[34] Building on these established mechanisms, future work should focus on elucidating how sustained Cu^2+^ release from PBGCu specifically influences these pathways over time, particularly under pathological conditions such as metabolic syndrome. For example, it remains unclear how prolonged copper exposure modulates macrophage polarization, redox balance, and endothelial cell function in impaired healing environments. In addition, the synergistic effects between bioactive glass-derived ions (e.g., Si^4+^, Ca^2+^) and Cu^2+^ in regulating cellular responses and extracellular matrix remodeling warrant further investigation. From a translational perspective, further studies are needed to evaluate the long-term safety and clinical efficacy of PBGCu formulations in treating infected wounds, with particular attention to potential cumulative copper toxicity, optimal dosing, and treatment duration. Furthermore, comparative studies against clinically used topical formulations would help establish the relative advantages of PBGCu and support its progression toward clinical application.

## 5. Conclusion

This study demonstrates the significant therapeutic potential of PBGCu formulations, particularly those containing 3 wt% copper, in enhancing wound healing and tissue regeneration in diabetes and metabolic syndrome. Both *in vitro* and *in vivo* findings support the efficacy of PBGCu in promoting epithelial regeneration, dermal tissue formation, and vascularization, while also exhibiting strong antibacterial properties. The controlled release of Cu^2+^ ions from PBGCu ointments provides sustained therapeutic benefits, making them particularly suitable for chronic wound care, especially in conditions like metabolic syndrome, where healing is often compromised. The ability of PBGCu to reduce scar formation and promote tissue remodeling further enhances its potential for clinical applications. Overall, these results highlight the promise of PBGCu as an effective wound healing agent, with future research focused on elucidating the molecular mechanisms underlying its effects and evaluating its safety and therapeutic efficacy in infected wounds.

## Acknowledgements

This work made use of the EPIC facility (RRID: SCR_026361) of Northwestern University’s NUANCE Center, which has received support from the SHyNE Resource (NSF ECCS-2025633), the IIN, and Northwestern’s MRSEC program (NSF DMR-2308691). This work is also supported by Querrey Simpson Institute for Regenerative Engineering at Northwestern University.

## Notes

### Competing Interest Statement

The authors have declared no competing interest.

