## Supplementary information for "A Pro-Regenerative Petroleum Jelly-Based, Copper-Doped Bioactive Glass Ointment for Impaired Wound Healing in Metabolic Syndrome"

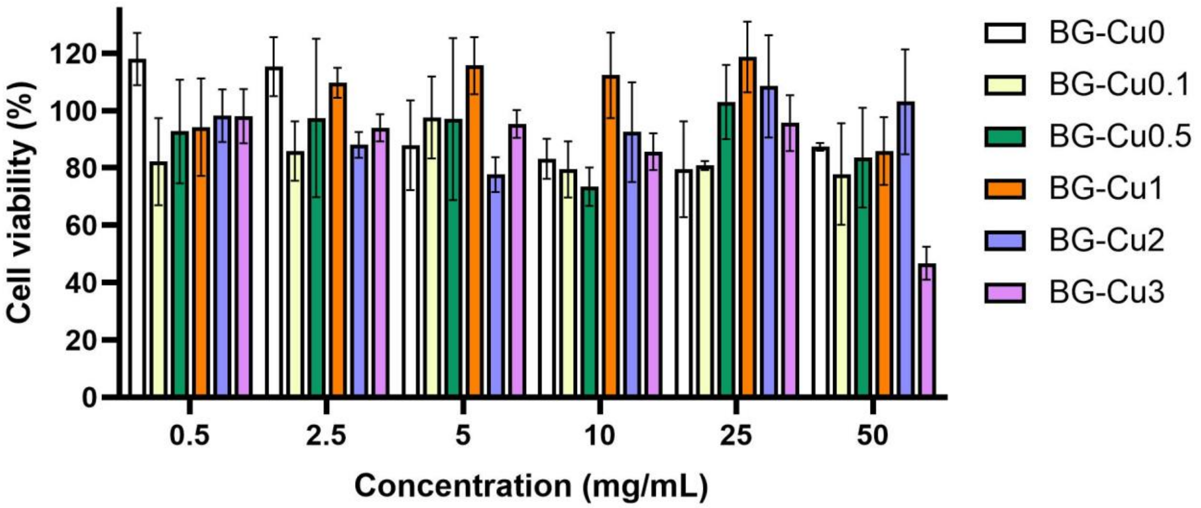


**Figure S1. Cell viability of human dermal fibroblasts after exposure to extracts from copper-doped bioactive glass powders (BG-Cu0 to BG-Cu3) at varying concentrations (0.5–50 mg/mL).** Viability was assessed using the MTT assay and is presented as a percentage relative to untreated controls. Data represent mean ± standard deviation (n = 3). All BG-Cu formulations maintained over 80% cell viability at concentrations up to 25 mg/mL.

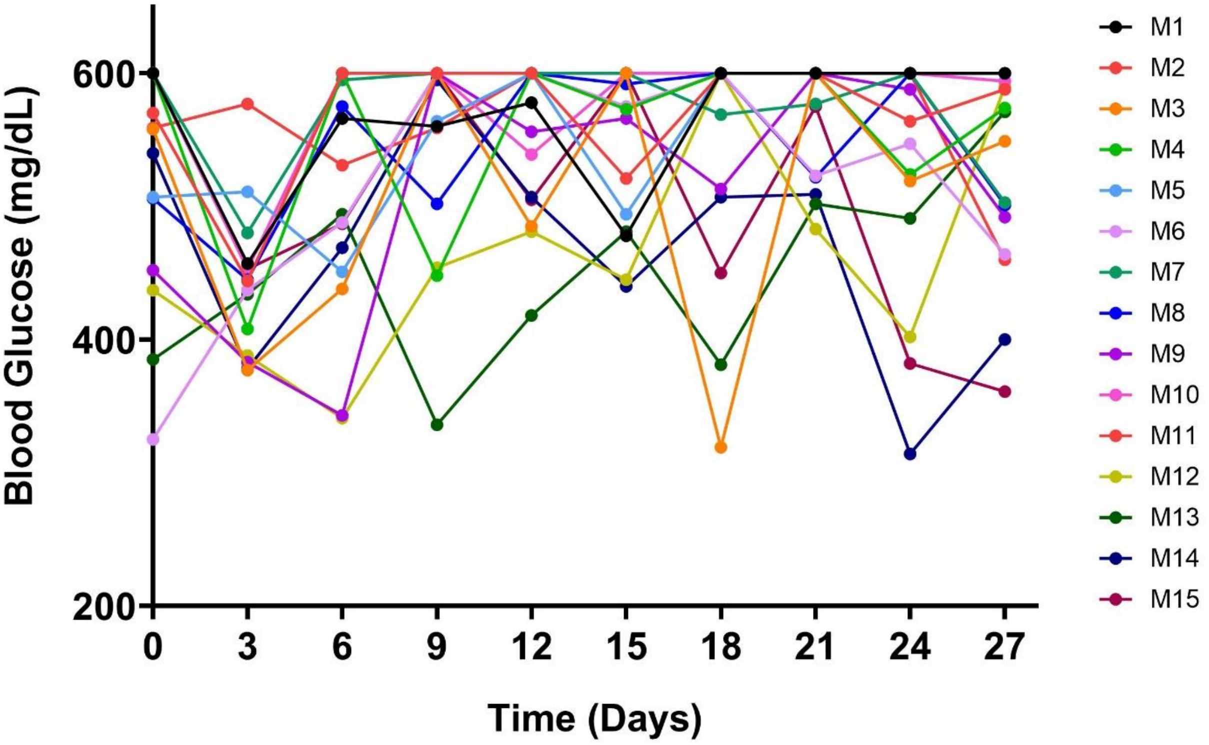


**Figure S2. Blood glucose levels in mice throughout the experimental period.** At all measured time points, blood glucose levels remained above 200 mg/dL in all mice.

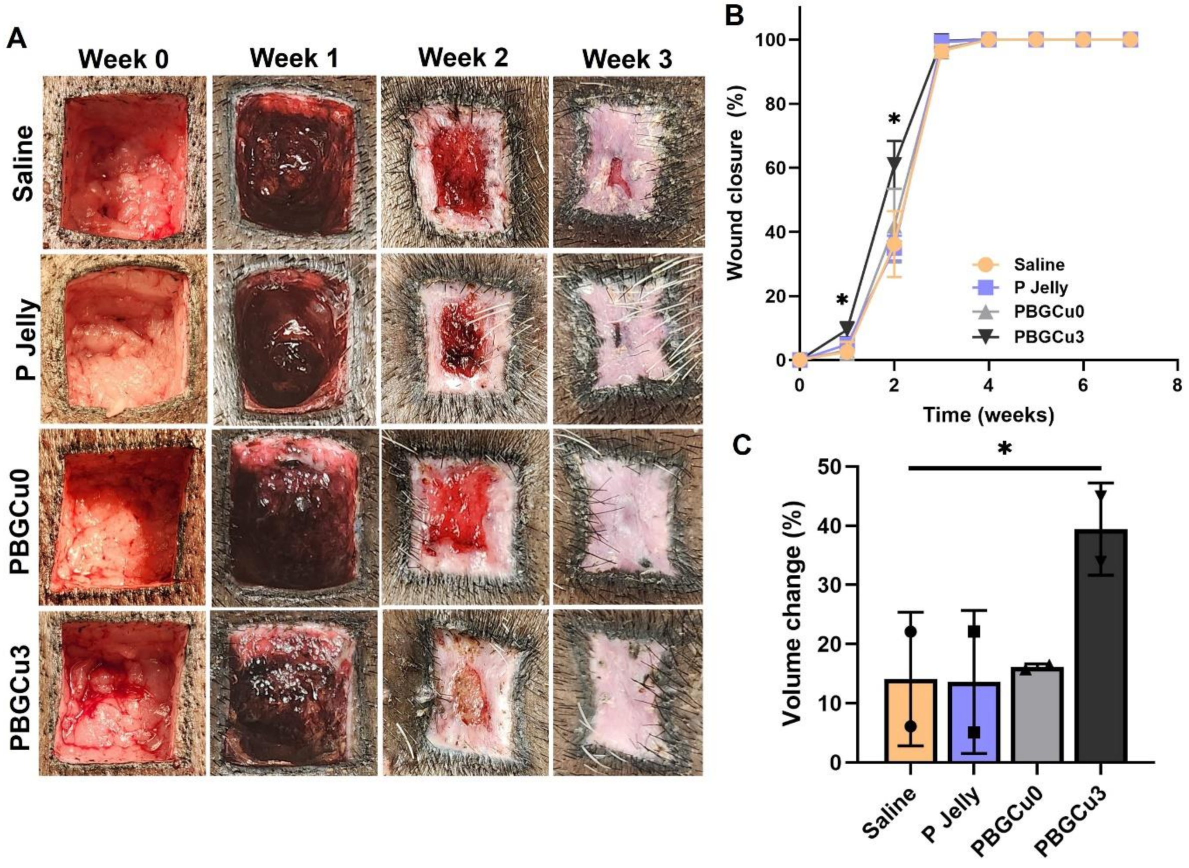


**Figure S3. PBGCu accelerates wound healing of full-thickness excisional wound in pigs with metabolic syndrome.** (**A**) Representative images showing wound closure at various time points (1, 2, 3, 4, 6, and 8 weeks) for each treatment group in Pig 2. (**C**) Quantification of wound closure rates over time in Pig 2, with PBGCu-treated groups showing significantly faster healing, and PBGCu3 demonstrating the most rapid closure. (**D**) Quantification of wound volume change after 1 week in Pig 2, with significant differences observed between PBGCu-treated groups and controls. All data are presented as mean ± standard deviation (SD), and statistical analysis was performed using one-way ANOVA followed by Bonferroni post hoc correction. ($n$ = 6, $^{**}p < 0.01$, $^{***}p < 0.001$, $^{****}p < 0.0001$).

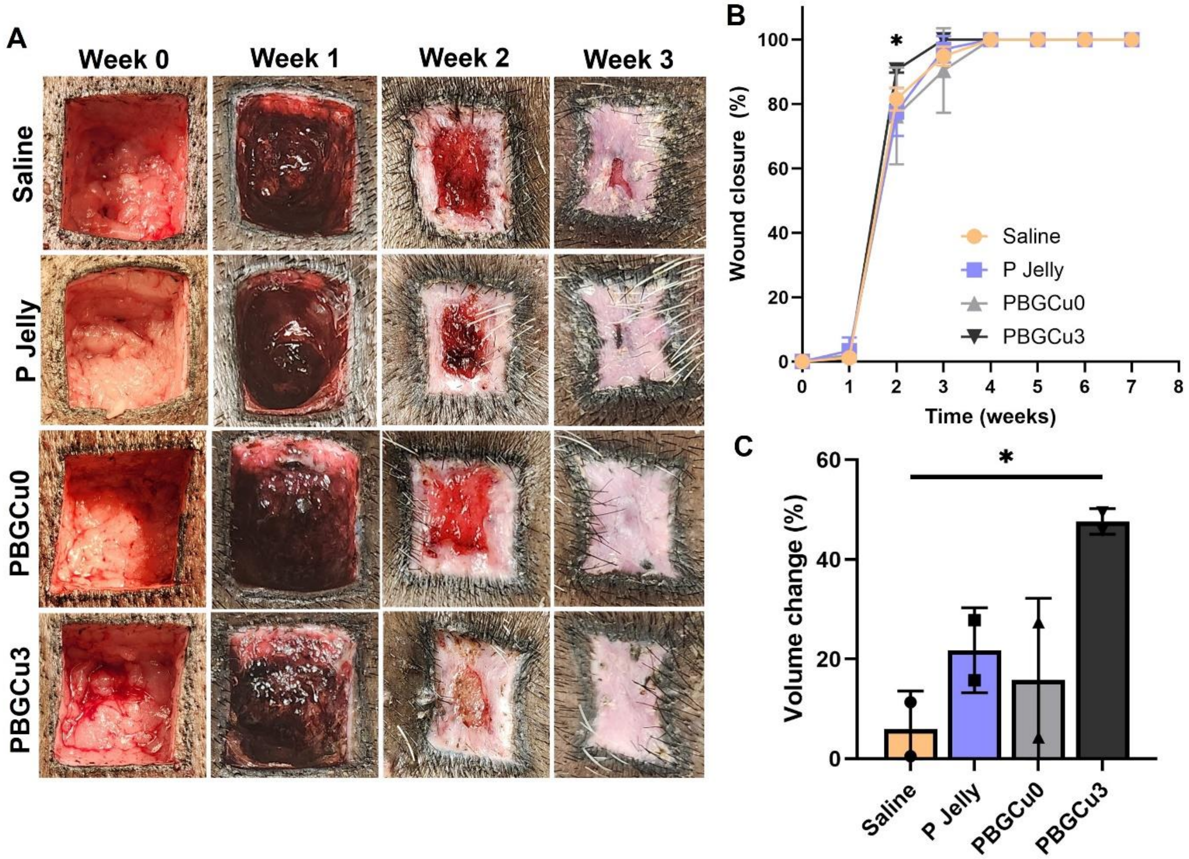


**Figure S4. PBGCu accelerates wound healing of full-thickness excisional wound in pigs with metabolic syndrome.** (**A**) Representative images showing wound closure at various time points (1, 2, 3, 4, 6, and 8 weeks) for each treatment group in Pig 3. (**C**) Quantification of wound closure rates over time in Pig 3, with PBGCu-treated groups showing significantly faster healing, and PBGCu3 demonstrating the most rapid closure. (**D**) Quantification of wound volume change after 1 week in Pig 3, with significant differences observed between PBGCu-treated groups and controls. All data are presented as mean ± standard deviation (SD), and statistical analysis was performed using one-way ANOVA followed by Bonferroni post hoc correction. ($n$ = 6, **p < 0.01, ***p < 0.001, ****p < 0.0001).

| Pig number | Cage number | Body weight (kg) | Blood glucose (mg/dL) | Cholesterol (mg/dL) | Triglyceride (mg/dL) |
|---|---|---|---|---|---|
| Pig 1 | 3386 | 75 | 102 | 269 | 116 |
| Pig 2 | 3411 | 100 | 52 | 392 | 77 |
| Pig 3 | 3385 | 97 | 67 | 263 | 18 |

**Table S1.** Blood test results for confirmation of metabolic syndrome phenotype in pigs.
